# VarFish - Collaborative and Comprehensive Variant Analysis for Diagnosis and Research

**DOI:** 10.1101/2020.01.27.921965

**Authors:** Manuel Holtgrewe, Oliver Stolpe, Mikko Nieminen, Stefan Mundlos, Alexej Knaus, Uwe Kornak, Dominik Seelow, Lara Segebrecht, Malte Spielmann, Björn Fischer-Zirnsak, Felix Boschann, Ute Scholl, Nadja Ehmke, Dieter Beule

## Abstract

VarFish is a user-friendly web application for the quality control, filtering, prioritization, analysis, and user-based annotation of panel and exome variant data for rare disease genetics. It is capable of processing variant call files with single or multiple samples. The variants are automatically annotated with population frequencies, molecular impact, and presence in databases such as ClinVar. Further, it provides support for pathogenicity scores including CADD, MutationTaster, and phenotypic similarity scores. Users can filter variants based on these annotations and presumed inheritance pattern and sort the results by these scores. Filtered variants are listed with their annotations and many useful link-outs to genome browsers, other gene/variant data portals, and external tools for variant assessment. VarFish allows user to create their own annotations including support for variant assessment following ACMG-AMP guidelines. In close collaboration with medical practitioners, VarFish was designed for variant analysis and prioritization in diagnostic and research settings as described in the software’s extensive manual. The user interface has been optimized for supporting these protocols. Users can install VarFish on their own in-house servers where it provides additional lab notebook features for collaborative analysis and allows re-analysis of cases, e.g., after update of genotype or phenotype databases.

## INTRODUCTION

Targeted sequencing (1) such as gene panel or whole exome sequencing (WES) has become common in clinical genetics research and diagnostic applications. Whole genome sequencing (WGS) is an emerging approach for such applications, yet interpretation of small non-coding variants remains challenging, and WES is still considered the most cost-efficient (2). Of course, the exome-like variants from WGS data can be analyzed in the same fashion as WES data. The interest in this area is shown by the large number of tools available for the scoring, filtering, and prioritization of exome-wide variants including Phen-Gen (3), OVA (4), BiERapp (5), QueryOR (6), wANNOVAR (7), MutationDistiller (8), eXtasy (9), or the Exomiser (10). These prior works cover different feature sets (shown in Table 1) and differ in stability and availability of their source code. The latter is particularly important when authors discontinue their service. Common features include variant pathogenicity and gene-phenotype similarity scores, annotations and filtering by population frequencies. In the light of database updates, we view certain topics to be important emerging themes in the field of variant filtering and prioritizing. These include joint filtering of multiple cases, collaboration in analysis, building databases of analysed cases with variant assessments, and the re-evaluation of cases. Approaching these topics commonly requires duplicate work or advanced bioinformatics skills. Here, we report on our web-based application VarFish developed to tackle these challenges. VarFish is freely available without login at https://varfish-kiosk.bihealth.org and a demo version showcasing features available on custom install is available at https://varfish-demo.bihealth.org. Its source code is available under the permissive MIT license at https://github.com/bihealth/varfish-server.

**Table 1.** Feature comparison of state-of-the-art tools for variant filtering and prioritization.

|  | Exo<br>mis<br>er | Phe<br>n-Ge<br>n | eXta<br>sy | OVA | Que<br>ry<br>OR | w-A<br>NN<br>OV<br>AR | BiE<br>Ra<br>pp | Mut<br>ati<br>on<br>Dis<br>tiller | VarF<br>ish |
| --- | --- | --- | --- | --- | --- | --- | --- | --- | --- |
| latest update | 2019 | 2014 | 2013 | 2015 | 2017 | 2017 | 2017 | 2020 | 2020 |
| HPO | ✓ | ✓ | ✓ | ✓ | ✓ | ✓ | ✓ | ✓ | ✓ |
| OMIM |  |  |  | ✓ | ✓ | ✓ |  | ✓ | ✓ |
| gene lists/regions | ✓ |  |  | ✓ | ✓ |  | ✓ | ✓ | ✓ |
| no registration | ✓ | ✓ | ✓ | ✓ |  | ✓ |  | ✓ | ✓ |
| gene info |  |  |  | ✓ | ✓ |  | ✓ | ✓ | ✓ |
| demo / tutorial |  | ✓ |  | ✓ | ✓ | ✓ | ✓ | ✓ | ✓ |
| dynamic report |  | ✓ |  |  | ✓ |  | ✓ | ✓ | ✓ |
| tabular download |  | ✓ | ✓ | ✓ | ✓ | ✓ | ✓ | ✓ | ✓ |
| multi-sample VCF | (✓) | ✓ |  |  | ✓ | (✓) | ✓ |  | ✓ |
| bring your server | ✓ | ✓ | ✓ |  |  |  | ✓ |  | ✓ |
| user annotations |  |  |  |  |  |  |  |  | ✓ |
| ACMG-AMP<br>class |  |  |  |  |  |  |  |  | ✓ |
| quality control |  |  |  |  |  |  |  |  | ✓ |
| multiple VCFs |  |  |  |  |  |  |  |  | ✓* |
| in-house DB |  |  |  |  |  |  |  |  | ✓* |
| collaboration |  |  |  |  |  |  |  |  | ✓* |

A tick mark (✓) indicates that the feature is present. “Dynamic reports” allows users to interactively change filter and sorting options. “Tabular downloads” allows users to download a spreadsheet (e.g. Excel) file with their results. “User annotations” allow users to leave flag, color coded, or free-text comments, “ACMG-AMPs support” assists users in creating variant assessments following the ACMG guidelines. “Quality control” allows users to perform quality control, e.g., for depth of coverage or relatedness of individuals. “Multi-sample VCFs” supports cases with more than one sample while “multiple VCFs” supports querying multiple cases at the same time. “Bring your server” allows users to create their own installations on their own server. “In-house DB” allows to build a database of variants identified at the user’s institution. An asterisk (\*) indicates that the feature is only available on installations on the user’s own server. Parentheses around the tick mark in “multi-sample VCF” row indicates that filtering is restricted to predefined models of inheritance.

## RESULTS

### Feature comparison with State-of-the-Art Tools

Table 1 shows the features implemented in VarFish in comparison to state-of-the art web tools (based on Figure 1 from (8)). The year of latest update is important as databases on variants and genes are growing quickly. The support of the Human Phenotype Ontology (HPO) and Online Mendelian Inheritance in Man (OMIM) is relevant for prioritizing single exomes. Filtering of genes and regions is of interest when characterizing patient cohorts for variants in known disease-causing genes. Furthermore, support for multi-sample files or even multiple VCF files at once allows more advanced analyses. Implementing quality control features is important to gauge the quality of data in an integrated platform. Dynamic variant reports greatly improve usability over repeated submission of queries to batch systems. Tabular downloads (e.g., as spreadsheet files) make it possible to archive cases on the user’s computer and store them in clinical or laboratory information management system. Supporting users in annotating variants with flags, colour codes extends such systems in the manner of a laboratory notebook. Supporting users in the ACMG-AMP (11) classification of variants is useful for creating diagnostic reports. The possibility to organize cases in projects with project-based access control further fosters collaboration in data analysis and allows for building in-house databases. Custom installations on the user’s own server can address data privacy issues. Finally, the availability of extensive documentation, tutorials and providing the tools without requiring user registration lowers the entry barrier.

**Figure 1.**
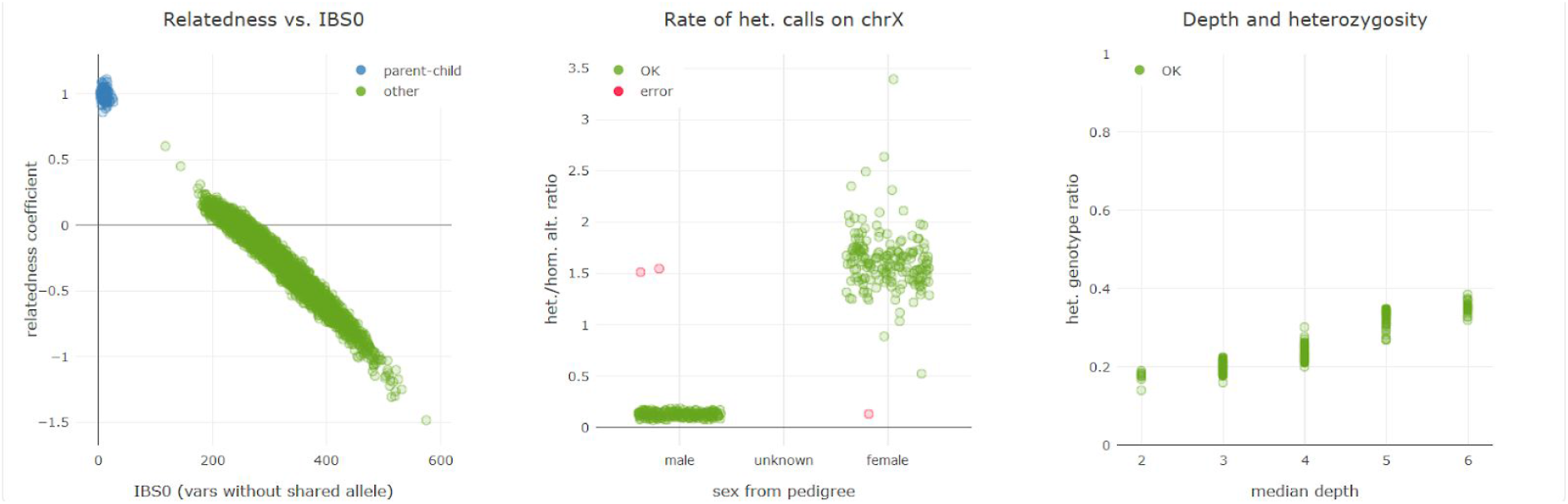
Quality control plots following the Peddy (22) approach. The plots are described in the main text.

### The VarFish Workflow

There are two major parts in the VarFish data processing workflow: the data *preprocessing and import* and the *query construction and execution* step.

The input to the *preprocessing and import* step is a file in VCF (Variant Call Format) format (12) and optionally a PLINK (13) pedigree file. Each variant record is read from the file and annotated with molecular impact using the Jannovar (14) library for both RefSeq (15) and ENSEMBL (16) transcripts, with the distance to the closest exon in either database, with its population frequencies in the ExAC (17), gnomAD (18), and Thousand Genomes Project (19) databases, and its presence in ClinVar (20). The variants are assigned a case identifier and are then imported together with properties such as quality scores from the VCF file into a PostgreSQL database table following the star schema pattern common in data warehouse applications.

The input to the *query construction and execution* step consists of the identifier of the case and the query settings from the user (see below). It constructs a SQL database query for selecting the variants based on the input criteria and joins the central variant table to further metadata tables (e.g., providing information about genes, variants, or conservation information). This query is then submitted to the database system for execution. VarFish was developed for use in genetics of rare diseases where users desire to create short lists of variants (say less than 200) for further analysis based on population frequency, genotype/segregation in families, molecular impact, and other criteria (21). In particular, the first three criteria can be used to greatly reduce the number of resulting variants. By employing the star schema pattern, database indices can be created for the most common queries and the (small) number of rows returned from the query on the central variant table can be obtained fast. The extensive metadata acquisition can then be limited to this small number of rows. As a periodic background job, VarFish tabulates the number of samples that each variant occurs in heterozygous, hemizygous, and homozygous state. This allows for removing variants seen in many cases as is the case for local polymorphisms or artifacts not seen in the population databases because of differences in variant calling.

### Quality Control Functionality

Another feature in VarFish is a global quality control function. Figure 1 shows an example of the quality control (QC) plots available. The three plots follow the Peddy methodology (22) and allow samples be examined for (un)expected relatedness, the sex derived from X-chromosomal variants, and depth-of-coverage vs. fraction of heterozygous variants. A detailed description and interpretation guide is available in the VarFish user manual. Further, VarFish allows users to provide quality control information for each sample that cannot be derived from VCF files, such as coverage information in JSON format (https://json.org). This allows an integrated display of QC information suitable for clinicians as suggested by Shyr, c.f. (23). Finally, the user can consider this information for all samples in a project to evaluate a sample in comparison to similarly processed ones or for a whole cohort.

### Database- and User-Based Annotation

VarFish integrates a growing list of databases. Databases such as gnomAD provide information on the variant frequencies within (sub)populations, dbSNP provides identifiers for registered variants, ClinVar provides variant pathogenicity and PubMed identifiers. Protein-level conservation information is derived from UCSC genome browser (24) data, the NCBI gene database provides gene summaries and gene reference into function (RIF) information, and the HPO (25) provides phenotype information. The background databases need to be imported when installing VarFish locally. An archive file with this data is provided for download. We provide the full Snakemake (26) workflow for downloading the data from open and free sources for reproducibility.

The VarFish result display lists extensive links to databases and data portals providing additional information. Furthermore, functionality for remote control of the integrative genome viewer (IGV) (27) and assessment of variants by tools such as MutationTaster (28) are provided. Resulting variant lists can be directly uploaded into MutationDistiller (8) for a complementary analysis. Further, users can annotate variants with flags, color codes, and free-form text as shown in Figure 2. This figure also shows the support for computing and storing using the ACMG-AMP guidelines (11).

**Figure 2.**
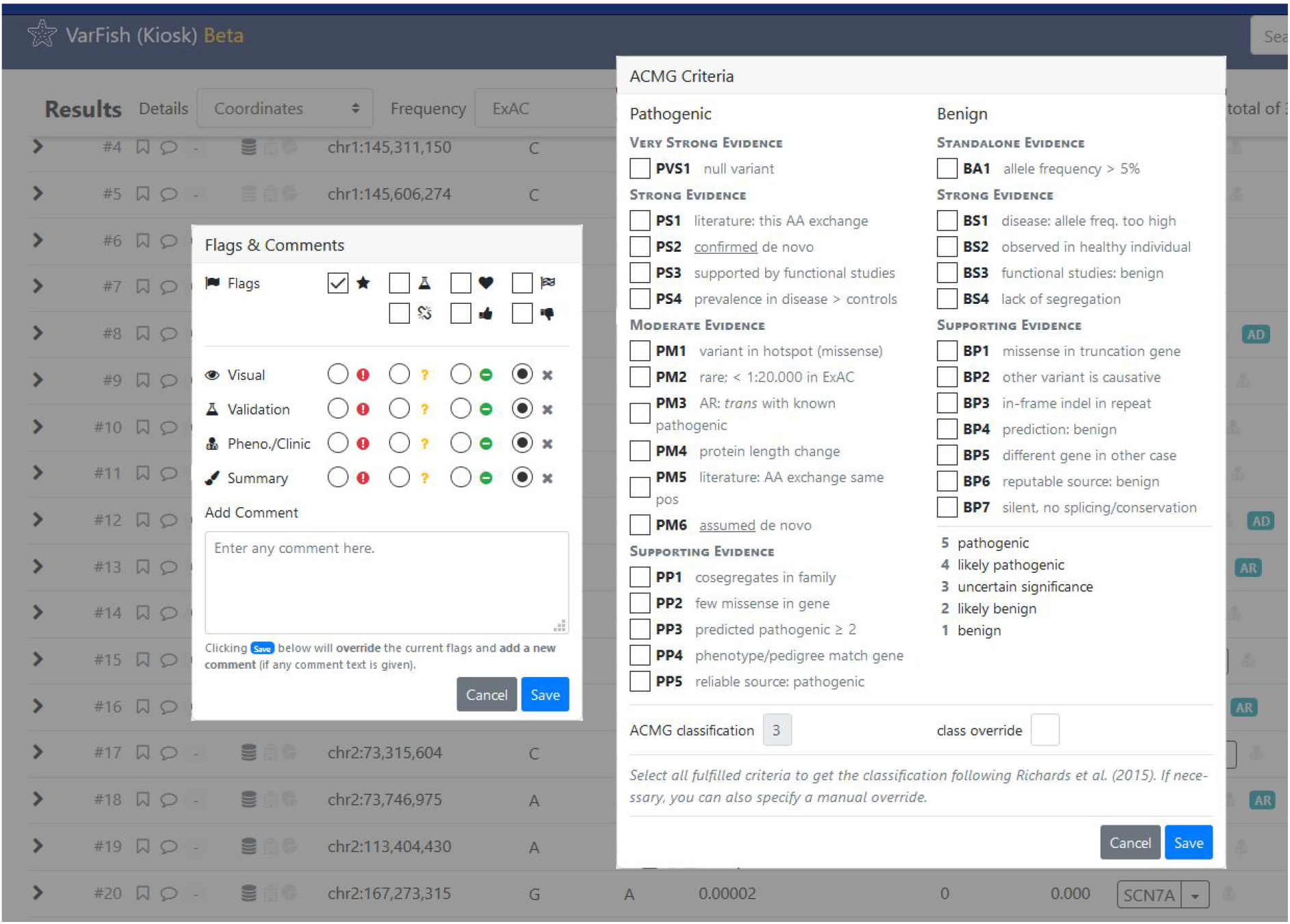
User annotation of variants. Users can apply flags and color codes to variants and leave free-text annotations. Flags include “bookmark”, “reported as candidate” and “final causative variant” as well as “no phenotype linked to gene”. Color codes can be assigned in categories “raw data visual inspection”, “gene clinical/phenotype match” and “validation results” as well as an overall summary color.

### Filtering Interface

Figure 3 illustrates the filtering interface and workflow from the user’s perspective. The aim is to provide an easy access for inexperienced users yet offer high flexibility for experts. This is realized by providing two levels of presets. With no preset, VarFish provides a high degree of configurability for genotype, population frequency, variant quality measures (including variant call quality and variant allelic balance), and ClinVar annotation. On the first preset level, it provides default settings for several categories. For example, there are separate “super strict”, “strict”, and “relaxed” population frequency settings under the assumption of dominant mode of inheritance, separate ones for assumed recessive mode of inheritance, and “strict” and “relaxed” settings for variant quality measures. On the second preset level, VarFish allows the user to select between settings such as “*de novo*”, “dominant inheritances”, and “recessive inheritance.” For example, the second-level preset “recessive inheritance” will set the genotype filter to return homozygous variants and compound heterozygous variants in the index (enforcing appropriate genotypes for the parents if present) with appropriate (yet strict) population frequencies and strict quality thresholds. The rationale is that users prefer to start reviewing a few promising variants first and then relax the filters while the total number of variants remains manageable. The user browses each variant in the raw data using IGV as well as the gene summary information and gene phenotype links. The results of this research can then be documented for each variant and flags are generated for the different categories (see above). In contrast, the second-level preset “*de novo*” sets the genotype filter to “heterozygous” for the index, and “reference” for all other members in the pedigree, the population frequencies are set to very restrictive values, while the quality thresholds are relaxed and deeply intronic variants with a distance of up to 100 bp are included. The rationale is that *de novo* variants are very rare in trios and that even non-coding and low-covered variants are worth getting inspected, the latter also under the aspect of possible mosaic disease causing variants (29).

**Figure 3.**
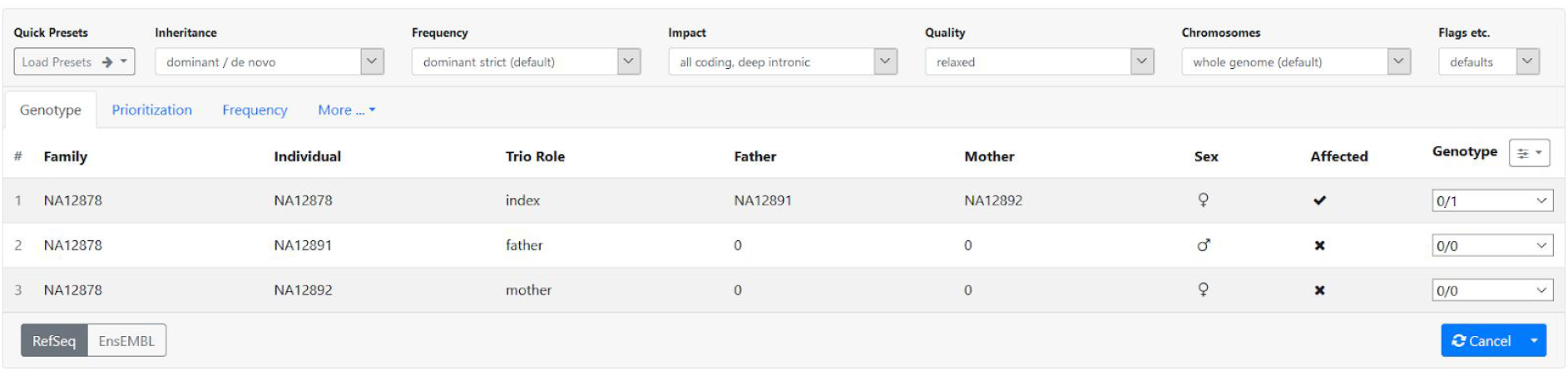
The filtering interface. XXX

### Joint Filtering of Multiple Cases

VarFish is capable of filtering the variants of multiple cases at once. The resulting variants are annotated with the number of cases that have at least one variant identified in a given gene. The result can then be sorted by that number to identify genes that carry rare variants of interesting impact and mode of inheritance. To demonstrate this, we performed a re-analysis of the original cohort used for identifying *TGDS* as a disease gene for Catel-Manzke syndrome published earlier (30). For this analysis, the second preset “recessive inheritance” was applied first, followed by relaxing the quality thresholds as the data was generated in the early days of WES sequencing. Figure 4 shows the top results. Of the resulting 388 variant records the only gene carrying variants in all six sequenced cases was *TGDS*, the next gene listed matched only two cases.

**Figure 4.**
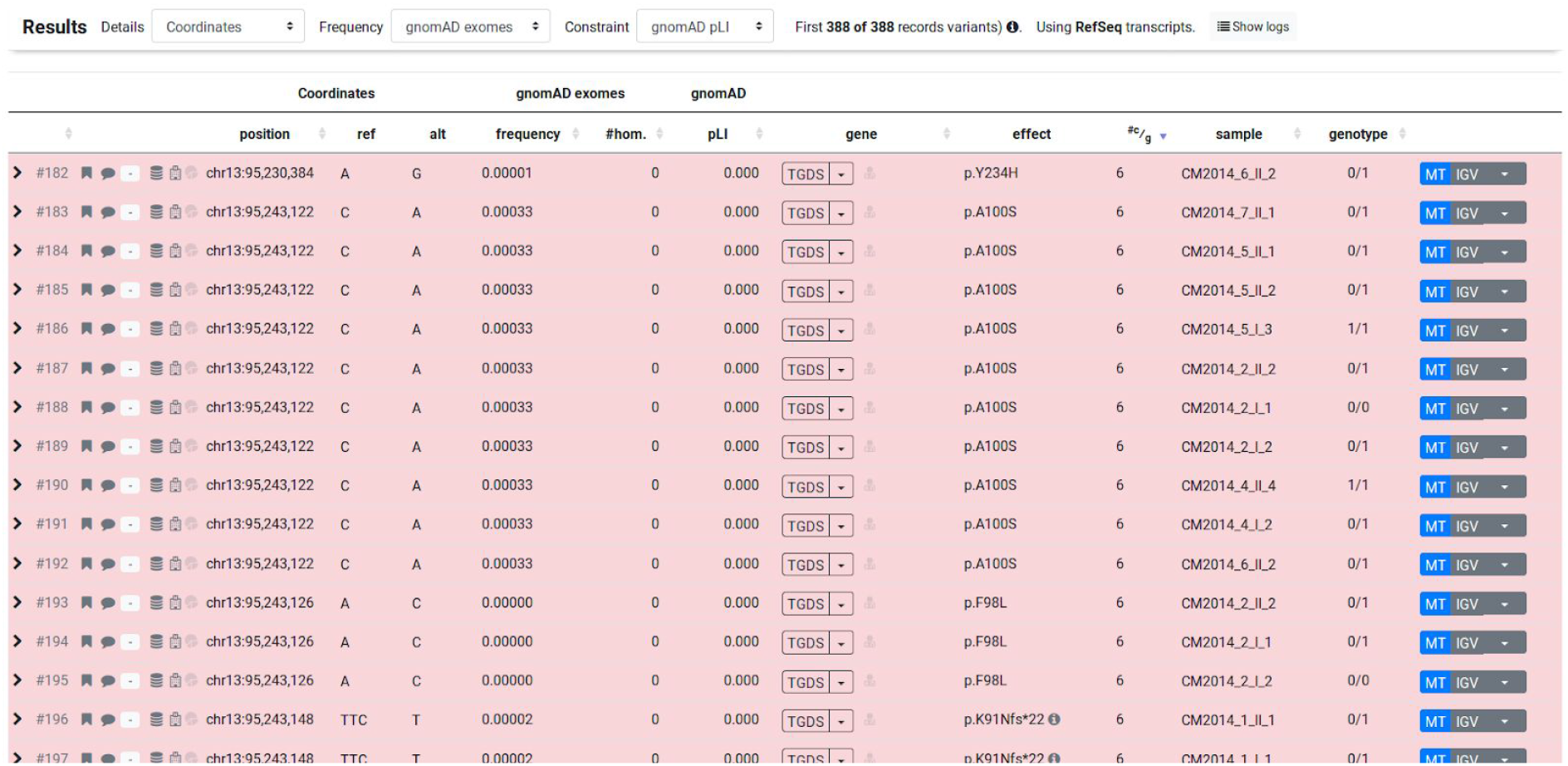
Catel-Manzke cohort filtering results (first 15 variants shown) to reproduce the finding that TGDS is the most likely candidate for being the disease gene.

## DISCUSSION

We here present VarFish, a flexible platform for the automated annotation of small variants, their filtering and prioritization. We demonstrated its use and effectiveness in a practical use case. The system aims at empowering biomedical and clinician researchers to perform complex, customizable variant prioritization and filtering in a maximally flexible way. Instead of implementing new custom scores, VarFish builds on and combines best-in-class scoring algorithms such as CADD and MutationTaster for variant pathogenicity prediction and gene-phenotype similarity computation. Annotation, filtering, and prioritization are shown to the user, allowing swift interactive processing by sorting variants according to different criteria. Comprehensive link-outs to external gene and variant databases are provided, and the integration of the IGV genome browser enables raw data inspection. Allowing the user to leave annotation flags, colour code, and free-text comments has proven highly useful for our in-house users. Data can be analyzed in an integrated platform up to ACMG-AMP variant assessment, followed by downloading a spreadsheet file for documentation in external systems.

Further advanced features such as collecting cases in projects and performing joint queries on multiple cases at once allow answering research questions that previously required bioinformatics expertise. For example, this allows an integrated characterization of disease cohorts by screening for pathogenic variants in known disease-associated genes followed by a joint analysis of the remaining cases, e.g., to identify jointly mutated genes.

Finally, the pipeline to generate background the database files from publicly available sources, and the pipeline for annotating VCF files and the filtering user interface is available under the permissive MIT open source license. This ensures full transparency, allows for setting up a fully reproducible variant analysis pipeline, and provides users with the advantage of open source systems in that there is no vendor-locking (in concordance with the FAIR (31) data management principles) and users are independent of the original software author. VarFish is used in the authors’ daily work and actively maintained. We welcome questions, comments, and suggestions via email or the Github project’s issue tracker.

## CONCLUSION

VarFish is a flexible and powerful platform for variant filtering, prioritization, and user-based annotation. With its collaboration and laboratory notebook features, it promotes collaborative analysis of cases and the re-analysis of variants at multiple points in time. Future development will focus on the extension of re-analysis features and supporting the analysis of structural variants, and support for whole genome data.

## WEB SERVER IMPLEMENTATION

VarFish is implemented in Python 3 with the Django web framework based on SODAR-core (https://github.com/bihealth/sodar_core) using PostgreSQL 11 for data storage and querying and VCFPy (32) for file parsing. In our installation, it runs on Linux container server with 128 GB of RAM, 16 cores, and 1 TB of disk.

## AVAILABILITY

VarFish is available for public usage at https://varfish-kiosk.bihealth.org. A demonstration instance with full collaboration features has been setup at https://varfish-demo.bihealth.org. The source code is available under the permissive MIT license from https://github.com/bihealth/varfish-server.

## ACCESSION NUMBERS

No new data was generated for this study. For demonstration purposes we used data from a previous publication (30).

## SUPPLEMENTARY DATA

Supplementary Data are available at NAR online.

## ACKNOWLEDGEMENT

The authors thank Hannah Kemmer, Jirko Kühnisch, and Elisa Schäfer for their feedback on VarFish.

